# The cost of data aggregation: spatiotemporal compression obscures causal attribution in biodiversity monitoring

**DOI:** 10.1101/2025.05.17.654658

**Authors:** Katarzyna Malinowska, Michał Wawrzynowicz, Katarzyna Markowska, Tomasz Chodkiewicz, Simon Butler, Lechosław Kuczyński

## Abstract

Conservation decision-making requires accurate identification of causes of population changes. Ecologists often rely on analytical protocols that aggregate high-dimensional monitoring data. We hypothesise that compressing data - either spatially, as in conventional time series (TS) analysis, or temporally, as in static species distribution models (SDMs) - destroys covariance structures and obscures the identification of causal drivers. To quantify this ‘aggregation cost’, we conducted a rigorous simulation experiment using virtual species to establish a known ‘ground truth’ of population drivers. We then employed a virtual ecologist approach to mimic a 20-year large-scale bird monitoring scheme, and generate realistic spatiotemporal datasets to evaluate the analytical pipelines. We benchmarked the causal attribution accuracy of aggregated TS and SDM protocols against a full-resolution spatiotemporal (FRST) framework, which retains native data dimensions and integrates mechanistic spatiotemporal covariance structures. Our simulations revealed that spatial compression severely compromises causal inference: unpenalised TS models failed to detect any true underlying drivers (accuracy = 0.50, sensitivity = 0.00). Temporal compression (SDMs) performed moderately better (accuracy = 0.68), while the FRST model achieved superior accuracy (0.88), sensitivity (0.84), and specificity (0.93). Furthermore, we identified a variable selection paradox: double penalty shrinkage marginally improved underpowered TS models, although it degraded the specificity of SDM and FRST frameworks by forcing spurious, correlated variables to absorb residual variance. Our findings demonstrate that protocols that involve data aggregation reduce the informational value of large-scale monitoring datasets. Full-resolution, mechanistically informed frameworks are essential for reliable causal attribution and robust biodiversity monitoring.

**Highlights:**

- Spatiotemporal data aggregation obscures causal drivers in biodiversity monitoring.
- Spatial compression in time series models fails to detect true population drivers.
- Full-resolution spatiotemporal models accurately identify true drivers.
- Automated variable selection introduces false positives in high-resolution models.

## 1. Introduction

Identifying the causes of population changes is a fundamental challenge in ecology. This task is especially relevant in the Anthropocene, when human-induced pressures accelerate both biodiversity declines (Ceballos et al., 2017; Díaz et al., 2019), and the expansion of invasive species (Nentwig et al., 2018; Seebens et al., 2021). In either context, whether reducing extinction risks or controlling biological invasions, effective management relies on the capacity to accurately attribute detected population changes to specific factors (Haubrock et al., 2023; Peery et al., 2004). This is essential for developing diagnostic indicators within monitoring frameworks, enabling a transition from passive observation to the development of conservation strategies.

A broad spectrum of analytical approaches can be used to identify factors underlying population change, from correlative methods to process-based, and more formal causal models (Dormann et al., 2012; Green, 1995; Peery et al., 2004; Schrodt et al., 2025). The latter have recently become more important in ecology as they provide mechanistic insight into observed patterns and enable causal inference (Grace, 2024; Schrodt et al., 2025). However, previous research on factors influencing population change has mainly relied on various correlative approaches (e.g., Smith et al., 2024; Spooner et al., 2018). Depending on the study and the purpose of the analysis, these approaches may involve compression of the spatiotemporal data. For example, they can be aggregated temporally into a single spatial unit (in time series analysis) or spatially in a single temporal snapshot (in species distribution models) (Erickson et al., 2017). Although the aggregation is a pragmatic solution to make the data statistically simpler, it could also discard the covariance structures inherent in the spatiotemporal data, potentially obscuring the true drivers of population dynamics (Johnson et al., 2024).

Standard time series analysis (TS) is used to examine trends and seasonal variations in observational data, without incorporating the spatial dimension (Moraffah et al., 2021; Shumway & Stooffer, 2016). Population size and the corresponding environmental data may be aggregated to the specific spatial scale at which the analysis is conducted, reducing the dataset to a single observation per time step. This compression creates a statistical bottleneck, as the effective sample size is limited strictly to the duration of the survey. Therefore, it could be challenging to make inferences, even with long-term time series, which are more difficult to obtain (Weisser et al., 2023; White, 2019). Furthermore, attempts to artificially boost effective sample sizes by combining heterogeneous time series replicates can actively degrade analytical accuracy, particularly when underlying inter-replicate variation is high (Slegers et al., 2025; Weisser et al., 2023). Ultimately, spatial compression may result in low statistical power and severely reduced ability to identify the true causes of population changes.

Species distribution models (SDMs) are a spatial counterpart to time series analysis. They have demonstrated remarkable versatility in predicting species distributions and understanding species-environment relationships (Dormann et al., 2012; Zurell et al., 2020). Typically, they combine environmental predictors with the species occurrence or abundance, without incorporating the temporal dimension, and treat populations as static snapshots (Elith & Leathwick, 2009; Guisan & Thuiller, 2005). SDMs are characterised by a number of limitations, such as the often-violated equilibrium assumption (Guisan & Thuiller, 2005; Zurell et al., 2009). Furthermore, a recent systematic review of two decades of SDM applications highlights a critical vulnerability: despite the rapid adoption of complex machine learning algorithms, the field is persistently hindered by oversimplified data structures, which severely limit robust ecological inference (Özkan Tümer et al., 2026). This issue is compounded by the fact that correlative SDMs rely on potentially biased sampling data, and subjective methodological decisions, which often lead to unreliable ecological forecasting (Barbosa & Alves-Souza, 2025; Jarnevich et al., 2015). Traditional SDMs usually ignore crucial ecological processes, such as density-dependent regulation, biotic interactions, or dispersal (Franklin, 2010; Holt, 2009; Normand et al., 2014), and have difficulty extrapolating models to a novel environment (Briscoe et al., 2019). They provide, however, a basis for the development of more advanced species distribution models that incorporate dynamic aspects such as phenology or demography (Zurell, 2017; Zurell et al., 2024). There is also a significant potential in demographic SDM or joint species distribution models (JSDM), which facilitate the modelling of species communities (Seoane et al., 2023; Zurell, 2017).

When spatiotemporal monitoring data are available, they are often aggregated across either spatial or temporal scales to fit traditional analytical methods. Therefore, we hypothesise that the data compression inherent in TS and SDM pipelines leads to high rates of false attribution due to severe information loss. In contrast, the full-resolution spatiotemporal (FRST) model, which retains the native resolution of the monitoring data, will correctly identify the drivers underlying population dynamics. To systematically evaluate this hypothesis, we conducted an extensive simulation experiment in which TS and SDM protocols were benchmarked against the proposed FRST model. We assessed their ability to correctly identify the true factors influencing population changes, by generating known ‘ground truth’. Because empirical datasets lack certainty regarding the underlying causal mechanisms, we used simulated ‘virtual species’ to provide an objective baseline with a fully known set of parameters for evaluating model performance (Hirzel et al., 2001). We subsequently applied the virtual ecologist framework (Zurell et al., 2010) to mimic the observational sampling process. While this method has been widely used to evaluate sampling biases and model predictive performance, its application to testing causal attribution remains largely limited.

## 2. Material and Methods

### 2.1 Study design

The primary objective of this study was to quantify the impact of spatiotemporal data aggregation on the accuracy of causal attribution in biodiversity monitoring. Because rigorous benchmarking of analytical protocols requires a known ground truth, we employed a ‘virtual ecologist’ framework (Malinowska et al., 2023). In the first phase of the study, we generated virtual species (VS) based on predefined relationships between environmental factors and population abundance. This allowed us to objectively evaluate the variable selection accuracy, and therefore the causal attribution performance, of each analytical method (Fig. 1). To ensure our simulations reflected true ecological and observational complexity, the virtual data were parameterised using real spatiotemporal abundance data from 10 western Palearctic passerine bird species (Table A.1). For each species, 100 virtual instances were generated, resulting in a total of 1000 independent replications. For the detailed algorithm of the study design see Appendix B.

**Figure 1.**
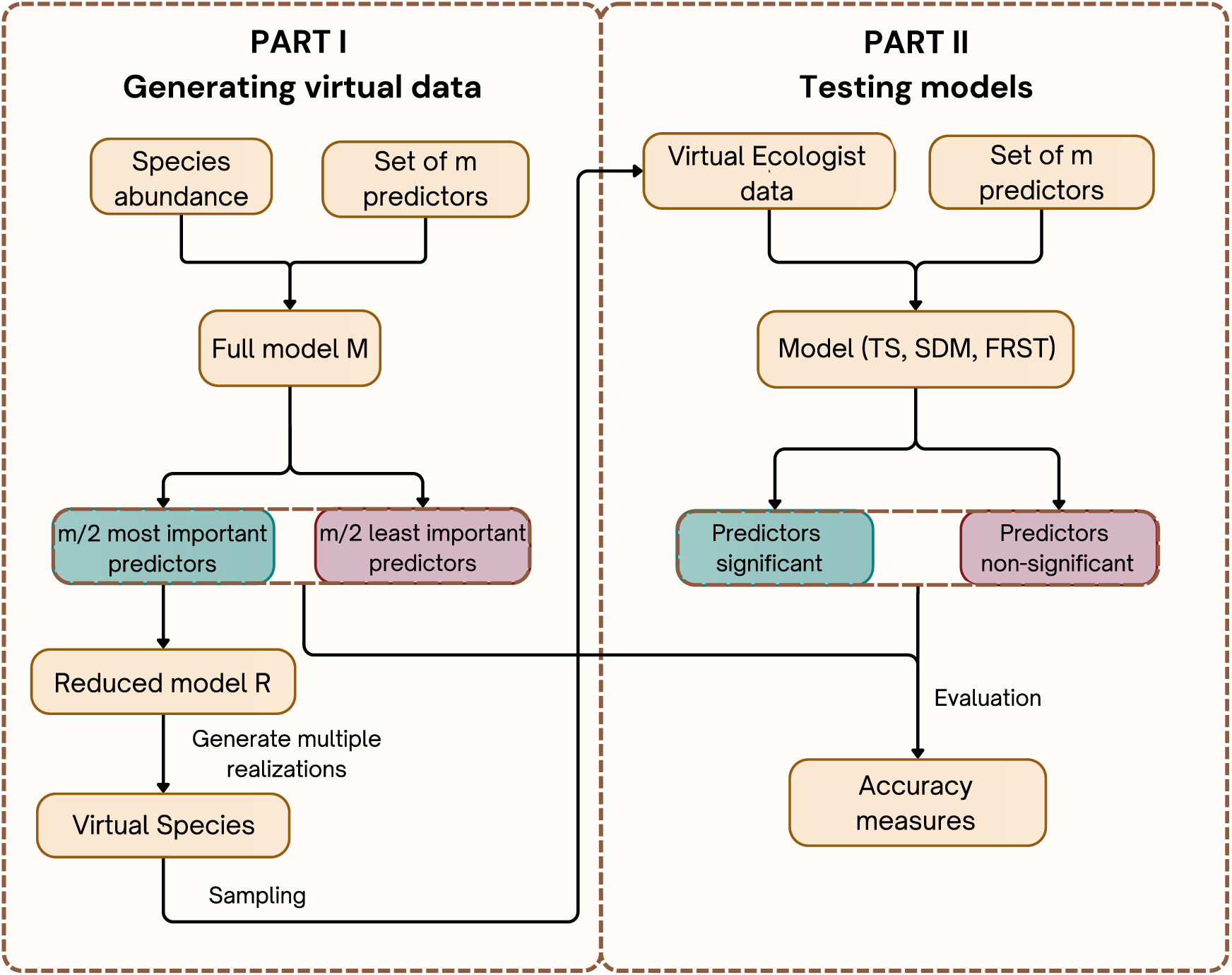
The study design. First, we fitted a model to the spatiotemporal bird abundance data using full set of predictors. We then selected half of the most important predictors from the full model to fit a reduced model. Multiple predictions were then generated from this reduced model, forming many instances of virtual species. By introducing a realistic level of randomness at this stage, each virtual species was different, even though it was generated from the same model. The sampling procedure, which mimicked data collection, produced virtual ecologist data. These data were then used as input to test the three analytical approaches – time series (TS), species distribution model (SDM), and the Full-Resolution Spatiotemporal (FRST) framework. Each of these models was fitted with the full set of predictors and then the variables were classified as “significant” using the p<0.05 criterion. As the true relevance of the predictors was known, it was possible to evaluate the three methods in terms of their effectiveness in identifying the real factors influencing population change.

### 2.2 Bird abundance data

To generate the virtual species, we used empirical abundance data from 14 common western Palearctic passerine birds, organised into 10 interacting pairs (meaning certain species were included in multiple pairings; see Table A.1 for details). These data were obtained from the Common Breeding Birds Survey in Poland (MPPL) and span a 20-year period (2001-2020). The MPPL monitoring programme strictly follows the methodology of the British Breeding Bird Survey (Gregory et al., 1996) and contributes to the Pan-European Common Bird Monitoring Scheme (PECBMS, Brlík et al., 2021). Specific details regarding fieldwork and data collection are presented in Appendix A. The MPPL programme provides raw counts, which were converted into species densities (pairs/km^2^) using standard distance sampling method (Buckland et al., 2001, 2004).

### 2.3 Environmental data

To parameterise the virtual environments, we extracted full-resolution spatiotemporal data for each MPPL survey site. This included dynamic, time-varying predictors such as annual land cover classifications derived from ESA CCI Land Cover (Defourny et al., 2017, 2024) and monthly climate variables from TerraClimate (Abatzoglou et al., 2018) (Table A.2). Because these abiotic variables are strictly bionomic - meaning they represent non-interacting, non-consumable conditions (Soberón & Nakamura, 2009) - we also incorporated a biotic interaction predictor to capture community-level dynamics (Wisz et al., 2013). Specifically, we modelled social information transfer, a process where individuals utilise heterospecific cues during habitat selection (Szymkowiak, 2013). To operationalise this mechanism, we included the dynamic abundance of an empirically validated interacting species as a biotic covariate. To correctly simulate heterospecific attraction and information use, this pool of candidate covariates was strictly limited to species with known positive interactions (Table A.1).

### 2.4 Virtual species and virtual ecologist

The virtual data, which serve as our known ‘ground truth’, were generated using the virtual ecologist framework described by Malinowska et al. (2023). The procedure can be summarised in four stages:

1. **Base model parameterization:** To ensure ecological realism, we initially fitted a Generalised Additive Mixed Model (GAMM; Wood, 2017) to the empirical data using the full set of 20 environmental and biotic covariates. GAMMs naturally accommodate non-linear species-environment relationships, zero-inflation, and complex hierarchical spatiotemporal structures. To establish a definitive ‘truth’ for causal attribution, we applied backward elimination to remove half of the covariates. The resulting reduced model, containing only the retained ‘causal’ drivers, was used as the blueprint to generate the virtual species. This explicit separation of causal from non-causal factors is the foundation of our variable selection tests.
2. **Virtual Species (VS) generation:** Predictions from the reduced model were computed to represent true virtual species abundance. To preserve the variance required for rigorous statistical testing, we introduced stochasticity by assigning random intercepts to sites and drawing random realisations from the original response distribution (Malinowska et al., 2023). This produced a fully resolved, spatiotemporal ‘true’ abundance for every site *i* and year *t*.
3. **Virtual Ecologist (VE) sampling:** To mimic the imperfections of actual monitoring, we simulated an observational process. For each virtual observer, a sampling error multiplier (*p*) was drawn from a lognormal distribution with a mean of 1 and a variance derived from the observer random effects estimated in the initial GAMM. The VE data were than obtained by multiplying the true VS densities by *p*.
4. **Calibration:** Uncalibrated VE data tend to overestimate prevalence and underestimate variance, therefore we applied quantile mapping to align the frequency distribution of the virtual data with the empirical observations. This calibration ensures the virtual datasets possess distributional properties indistinguishable from real ecological data (Malinowska et al., 2023).

Steps 2-4 were repeated 100 times for each of the 10 species pairs, yielding a total of 1000 independent spatiotemporal datasets. These VE datasets served as the ground truth to benchmark the causal attribution accuracy of the evaluated analytical frameworks.

### 2.5 Statistical experiment

Ecological relationships, particularly species’ environmental preferences, are frequently non-monotonic or unimodal, therefore summarising these effects with a single parameter or effect size can be misleading. Consequently, our study evaluates analytical performance based on binary causal attribution: the ability of a protocol to correctly classify a predictor as “significant” or “insignificant” in driving population dynamics. To provide a flexible, standardised baseline across all tested frameworks, we utilised Generalised Additive Models (GAMs; Wood, 2017). GAMs naturally accommodate non-linear responses by penalising the smoothness of relationships, effectively reducing to generalised linear models (GLMs) if the data do not support complex functional forms.

#### 2.5.1 Spatial compression (Time series; TS)

To implement the standard TS protocol, we applied spatial compression to the virtual ecologist datasets. Both the simulated population densities and each of the 𝑚 environmental variables were averaged across all spatial sites for each year. A GAM was then fitted to the temporally varying, but spatially aggregated, data:

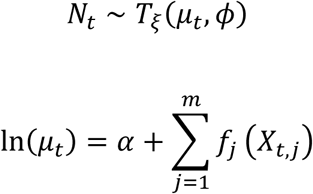

Here, population density 𝑁_𝑡_, assumed to follow a Tweedie distribution with index parameter 𝜉 and dispersion parameter 𝜙, is indexed solely by the year 𝑡. The natural logarithm of the expected population density is the sum of the intercept 𝛼 and smooth functions 𝑓_𝑗_, each fitted to the temporally varying predictor 𝑋_𝑡𝑗_.

#### 2.5.2. Temporal compression (Species distribution model; SDM)

To implement the static SDM protocol, we applied temporal compression. Population densities and predictors were averaged across the 20-year period to create a single static snapshot for each 1×1 km survey plot.

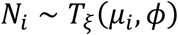

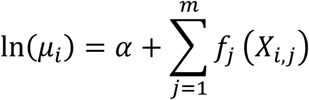

This structure mirrors the TS model, but the density 𝑁_𝑖_ varies exclusively in space, indexed by the sampling site identifiers 𝑖.

#### 2.5.3 Full-resolution spatiotemporal benchmark (FRST framework)

The FRST framework evaluates the data at its native resolution, retaining both spatial and temporal dimensions without aggregation. To appropriately model the covariance structures inherent in high-dimensional ecological data, this framework explicitly embeds mechanistic proxies: density-dependence, spatiotemporal autocorrelation, and hierarchical random effects. Density dependence was modelled using Gompertz dynamics. For any site 𝑖, the population growth rate 𝑟 = ln(𝑁_𝑡_/𝑁_𝑡−1_) implies ln(𝑁_𝑡_) = ln(𝑁_𝑡−1_) + 𝑟. Substituting 𝑟 with a smooth, non-parametric function 𝑓_𝑑_ allows for flexible self-regulation (including non-monotonic Allee effects):

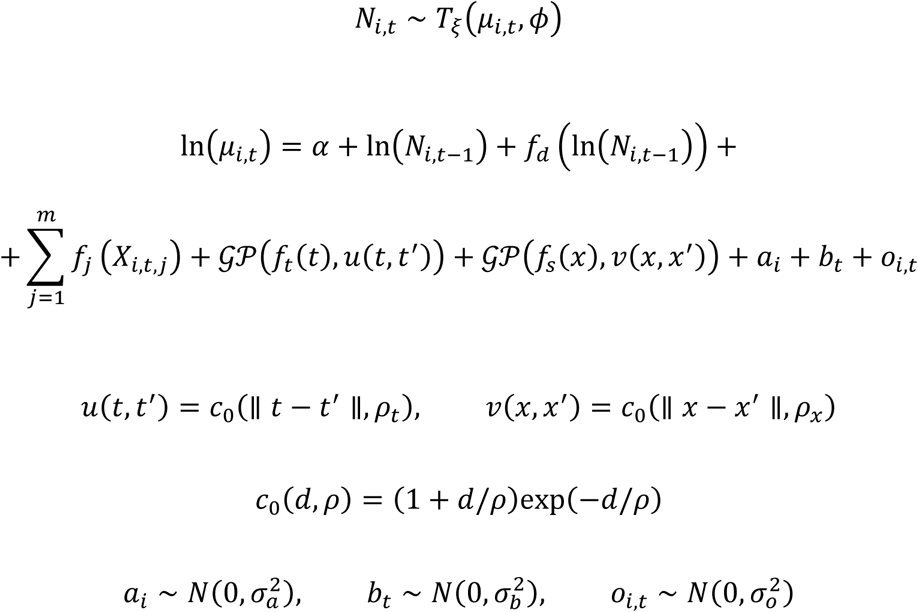

Unlike the aggregated models, all 𝑚 predictors (𝑋_𝑖,𝑡,𝑗_) vary dynamically in space and time. The structural extensions to handle this resolution include:

1. Temporal autocorrelation: Modelled as a Gaussian process (𝒢𝒫) using the Matérn correlation function 𝑐_0_ with parameter 𝜌_𝑡_ to describe between-year covariance.
2. Spatial autocorrelation: Modelled as a 𝒢𝒫 describing geographic covariance between sites using a Matérn function with parameter 𝜌_𝑥_.
3. Random spatial and temporal variation: Normally distributed random intercepts (𝑎_𝑖_, 𝑏_𝑖_) with the mean zero and variances σ*_a_*^2^,σ*_b_*^2^ to absorb unmeasured environmental noise.
4. Observational process: A normally distributed random intercept (𝑜_𝑖,𝑡_) with variance σ_0_^2^ to account for observer-specific detection biases.

### 2.6 Methods evaluation

To benchmark the causal attribution performance of the TS, SDM, and FRST frameworks, we evaluated their ability to correctly recover the underlying drivers from the 1000 virtual ecologist datasets. Because variable selection is a common strategy to improve model interpretability, all frameworks were fitted in two configurations: one without automated variable selection, and one utilizing double penalty shrinkage (Marra & Wood, 2011). While shrinkage approaches are well-documented to improve predictive accuracy by penalizing the size of smooth functions toward zero, their efficacy for causal attribution, particularly under severe spatiotemporal data aggregation, remains underexplored. Computationally, this shrinkage was invoked using the `select=TRUE` argument in the ‘`mgcv`’ package (Wood, 2017). For all configurations, a predictor was classified as ‘significant’ (selected) if its estimated p-value was < 0.05. While arbitrary, this threshold reflects the standard heuristic employed in applied ecological research and conservation reporting, ensuring our findings translate directly to practitioner workflows.

By comparing the variables retained by each model against the known ‘ground truth’ of the virtual species, we calculated three binary classification metrics:

1) Accuracy (correct classification rate): The overall proportion of correctly classified variables (i.e., true causal drivers successfully identified and spurious non-causal variables successfully excluded).
2) True positive rate (sensitivity): The power of a test, that is the probability that a true causal driver used to generate the virtual data was successfully detected by the model.
3) True negative rate (specificity): The probability that a non-causal, spurious variable not used to generate the virtual data was correctly ignored by the model.

To formally test the effects of data aggregation and selection strategy on these performance metrics, we fitted binomial Generalised Linear Mixed Model (GLMM) using the “glmmTMB” package (Brooks et al., 2017). The analytical framework (TS, SDM, FRST), the variable selection strategy (with/without shrinkage), and their interaction were included as fixed effects, while the virtual species was modelled as a random intercept to account for taxon-specific baseline differences. Pairwise contrasts were estimated via marginal means using the “emmeans” package (Searle et al., 1980). All analyses were performed in R 4.5 software (R Core Team, 2025). The R code used for the analysis and data associated with final results are available at: https://github.com/popecol/Causes.

## 3. Results

### 3.1 Accuracy

The spatially compressed Time Series (TS) model exhibited the lowest causal attribution accuracy overall (Figs 2a and B.1, Table B.1). Without automated variable selection, TS accuracy was effectively random at 0.50 (95% CI: 0.48–0.53). Applying double penalty shrinkage marginally improved this to 0.55 (CI: 0.52–0.57; Odds-Ratio (OR): 0.83, p<0.0001; Table B.2). The temporally compressed Species Distribution Models (SDMs) performed moderately better, achieving a mean accuracy of 0.68 (CI: 0.66–0.70) without shrinkage. However, contrary to the TS protocol, applying shrinkage to SDMs significantly reduced accuracy to 0.63 (CI: 0.61–0.65; OR: 1.24, p<0.0001). Under both configurations, SDMs were significantly more accurate than TS (OR: 0.47 and 0.71, p<0.0001; Table B.3). Retaining native data resolution yielded the best performance: the FRST framework significantly outperformed both aggregated approaches, achieving an accuracy of 0.88 (CI: 0.87–0.89) without shrinkage, dropping to 0.77 (CI: 0.76–0.79) when shrinkage was applied (OR: 2.22, p<0.0001). The FRST framework remained significantly more accurate than both SDM (OR: 0.28 and 0.50, p<0.0001) and TS (OR: 0.13 and 0.36, p<0.0001).

**Figure 2.**
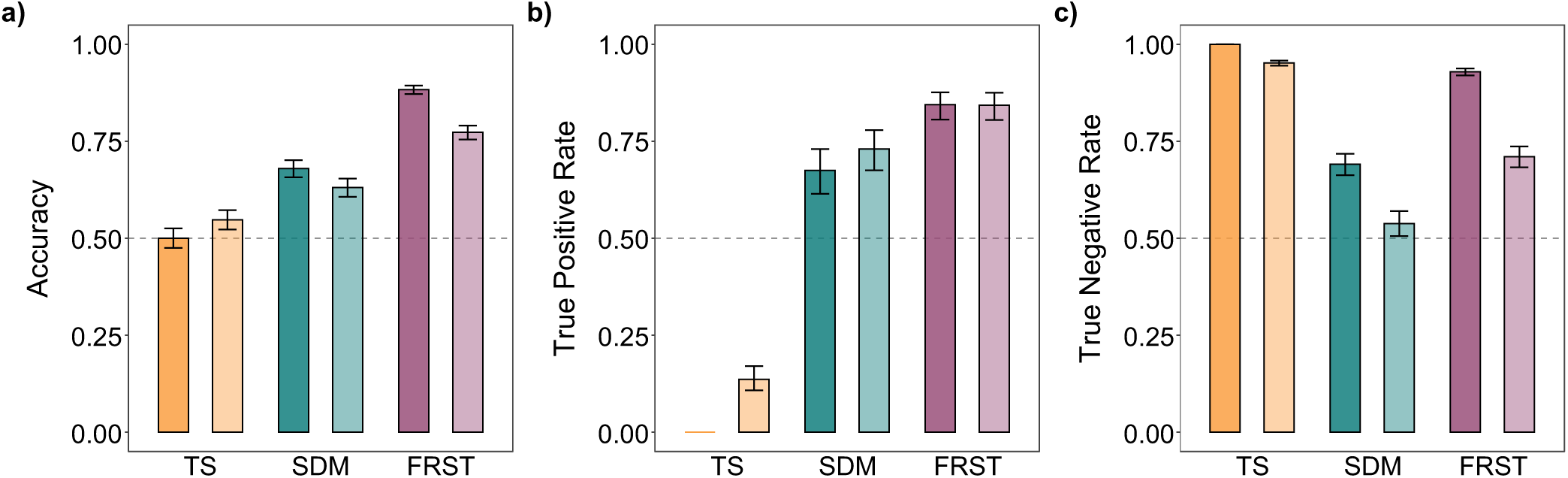
Performance metrics such as a) accuracy, b) true positive rate, and c) true negative rate of the time series (TS), species distribution model (SDM), and full-resolution spatiotemporal (FRST) approach in identifying population drivers. Column heights represent mean values and error bars indicate 95% confidence intervals. A darker column color within the approach indicates a lack of variable selection, while a lighter color indicates that double penalty shrinkage was used. The baseline at 1/2 indicates expected value if the variables were classified randomly.

### 3.2 Sensitivity

The cost of spatial aggregation was most evidently revealed in the models’ sensitivity (true positive rate). Without shrinkage, the TS approach demonstrated no statistical power, failing to detect any of the true causal drivers across all simulations (sensitivity = 0.00, CI: 0.00 – 1.00; Figs 2b and B.2, Table B.1). The enforcement of variable selection minimally enhance sensitivity to 0.14 (CI: 0.11–0.17). In contrast, temporally compressed SDMs successfully recovered a moderate proportion of true drivers (0.68, CI: 0.62–0.73), with shrinkage further improving sensitivity to 0.73 (CI: 0.68–0.78; OR: 0.77, p<0.0001; Table B.4). The FRST framework demonstrated the highest statistical power of all tested methods, identifying the vast majority of true causal drivers (0.84, CI: 0.81–0.88). For the FRST approach, the choice of variable selection strategy did not significantly alter sensitivity (OR: 1.0, p=0.79). FRST significantly outperformed SDM under both configurations (OR: 0.38 and 0.50, p<0.0001) and surpassed TS when variable selection was applied (OR: 0.03, p<0.0001; Table B.5).

### 3.3 Specificity

The TS approach without variable selection demonstrated an apparent high specificity at 1.00 (CI: 0.00 – 1.00; Figs 2c and B.3, Table B.1). However, this is a statistical artifact: since the aggregated model lacked the degrees of freedom to detect any variables (yielding zero sensitivity), it necessarily produced zero false positives. When double shrinkage forced the TS model to select variables, specificity dropped to 0.95 (CI: 0.95–0.96). SDMs struggled significantly with false positives, specificity was 0.69 (CI: 0.66–0.72) without shrinkage and dropped severely to 0.54 (CI: 0.51–0.57) when shrinkage was applied (OR: 2.00, p<0.0001; Table B.6). The FRST framework successfully maintained robust specificity at 0.93 (CI: 0.92–0.94) without shrinkage. However, applying automated variable selection introduced substantial false positives, significantly degrading FRST specificity to 0.71 (CI: 0.68–0.74; OR: 5.00, p<0.0001). Despite this drop, the baseline FRST model outperformed SDM in avoiding spurious correlations across all configurations (OR: 0.00, p<0.0001; Table B.7) and outperformed TS when selection was applied (OR: 8.0, p<0.0001). Kernel density plots of performance metrics for each modelling approach (Figs B.4 and B.5), as well as for each modelled species (Figs B.6 and B.7) confirm this consistent hierarchy of effectiveness among the analysed approaches in both variable selection variants.

## 4. Discussion

Our findings demonstrate the severe analytical cost of spatiotemporal data aggregation when inferring the causes of population change. By evaluating methods against a known virtual ground truth, we revealed that the time series (TS) approach, which involves spatially compressed data, systematically fails to detect factors influencing population abundance. In our unpenalised simulations, TS classified all variables as non-significant, regardless of their actual causal impact. In contrast, species distribution models (SDMs) that operate on temporally compressed data performed significantly better, achieving moderate accuracy (0.68) with a better balance of sensitivity and specificity. The full-resolution spatiotemporal (FRST) framework demonstrated the best overall performance, achieving the highest accuracy (0.88) and exceptional statistical power (0.84) by evaluating the monitoring data at its native resolution.

Naturally, the choice of analytical protocol is constrained by data availability and the specific research hypotheses. As Gould et al. (2025) demonstrated, a single research problem can provide a different set of answers depending on the modelling framework employed. For instance, time series models are the most effective tool for capturing interannual fluctuations in apparent survival, when analysing purely demographic data, as these often vary more temporally than spatially (Auger-Méthé et al., 2021; Salewski et al., 2013). Conversely, SDMs may be better suited to analyse large-scale environmental gradients, which typically exhibit stronger spatial than temporal structure (Franklin, 2010; Ortiz-Yusty et al., 2023). However, when the objective is to identify the drivers of population change using spatiotemporal abundance data, our results indicate that employing correlative approaches on aggregated data carries underappreciated risks.

### 4.1 The cost of spatiotemporal compression

Although TS and SDMs remain ‘first choice’ ecologists tools in conservation ecology, our results show that they provide misleading causal inferences. The main reason for their poor performance in our experiments is data aggregation. Spatial or temporal compression of data fundamentally alters its statistical properties: variance is collapsed, degrees of freedom are radically decreased, and the signal-to-noise ratio is artificially modified (Pollack et al., 2022; Pollet et al., 2015; Schad et al., 2024). In consequence the validity of statistical inference is reduced, and variable selection is severely biased (Abrahamowicz et al., 2004). Our results indicate that the greater the level of data aggregation, the more granularity is lost and the more severely true effect sizes are attenuated. This perfectly explains the statistical failure of TS (spatial compression), the moderate performance of SDM (temporal compression), and the high accuracy of the FRST framework, which preserves the original fine scale spatiotemporal patterns. Similar aggregation-induced biases have been detected in other time series analysis, where datasets averaged over inadequate temporal scales exhibit weakened covariate effects (Ferguson et al., 2017), and where the temporal aggregation of environmental variables fundamentally alters the performance and reliability of predictive ecological models (Salles et al., 2016).

### 4.2 Properties of the FRST framework

The superior performance of the FRST framework is driven by the use of unaggregated spatiotemporal data and its modelling structures, which are designed to handle associated data complexity. By explicitly incorporating mechanistic proxies of the data-generating process, such as Gompertz density dependence, spatiotemporal autocorrelation (Gaussian processes), and observer effects, the framework accurately partitions true environmental signals from structural noise. Ignoring this hierarchical structure typically leads to biased inferences, whereas mechanistically informed models provide a much more accurate reflection of ecological reality (Franklin, 2010; Zurell, 2017).

However, the FRST approach has also some limitations. Retaining the native resolution of large-scale monitoring data demands significant computational resources. The complexity of fitting spatiotemporal GAMMs dramatically increases simulation times, data preparation efforts, and the technical expertise required for model tractability, and its understanding. Moreover, spatiotemporal models require large amounts of data, which might be difficult to obtain. Whether this analytical investment is justified depends on the availability of high-resolution data and the need for diagnostic accuracy in the specific conservation problem.

### 4.3 The paradox of variable selection

Our study contributes to the ongoing debate regarding automated variable selection in ecology (Tredennick et al., 2021; Whittingham et al., 2006). We demonstrated that double penalty shrinkage only marginally improves the accuracy of fundamentally flawed TS models. More importantly, forcing automated variable selection actively reduced the efficiency of the more powerful SDM and FRST approaches by introducing substantial false positives (lowering specificity). This suggests that, when a model possesses sufficient degrees of freedom and is correctly specified structurally, aggressive shrinkage can worsen collinearity and concurvity issues in highly parameterised models. By heavily penalizing the smooth functions of true causal drivers, the algorithm may force spatially or temporally correlated spurious variables to inappropriately absorb the residual variance. This aligns with broader theoretical warnings that using statistical correlation methods to select variables in SDMs often results in highly unstable and inconsistent sets of covariates (Díaz-Vallejo et al., 2024). Furthermore, our conclusions are supported by recent virtual ecologist simulations demonstrating that regularised variable selection methods often underperform compared to simpler approaches in identifying true ecological drivers without pulling in spurious covariates (Cushman et al., 2024). This variance-shifting inflates the statistical significance of non-causal factors, thereby reducing specificity. As previously cautioned in the modelling literature, increasing the complexity of algorithms to handle high-dimensional data cannot substitute for robust a priori ecological justification. Doing so, frequently results in biologically spurious attributions (Bell & Schlaepfer, 2016). Consequently, for high-resolution spatiotemporal models, fitting the full model, driven by structural, mechanistic hypotheses rather than aggressive selection penalties, may yield the most reliable causal inferences.

### 4.4 Practical implications and conclusions

This study evaluates the effectiveness of analytical approaches used to determine the causes of population change against the known ground truth. Although we emphasize the critical importance of fine-grained, long-term, and large-scale monitoring data, such as that provided by the Pan-European Common Bird Monitoring Scheme (Brlík et al., 2021), our results demonstrate that the value of these datasets is lost if they are subsequently compressed to fit low-dimensional analytical protocols. We do not claim to have tested every available statistical tool used for identifying causes of population change, however, our findings clearly illustrate that standard data aggregation protocols can obscure true causal drivers and lead to erroneous management conclusions. By adopting accessible, full-resolution methods like the FRST framework, which account for both ecological mechanisms and observational realities, researchers can accurately translate modern monitoring data into reliable biodiversity indicators for mitigating the effects of global change.

## Data availability statement

The code and data associated with the final results are available on the GitHub repository: https://github.com/popecol/Causes. Bird population data used in the analysis are partially owned by OTOP BirdLife Poland (2000-2006) and Chief Inspectorate for Environmental Protection (GIOŚ, 2007-2021). OTOP provides the data upon request. GIOŚ provides the data at: https://monitoringptakow.gios.gov.pl/PM-GIS/?lang=en, without rights to redistribute them. Environmental data, such as climate data are publicly available at the page: https://www.climatologylab.org/terraclimate.html, as well as land cover data: https://maps.elie.ucl.ac.be/CCI/viewer/download.php.

## CRediT authorship contribution statement

**Katarzyna Malinowska:** Conceptualization, Data curation, Formal analysis, Methodology, Software, Visualization, Writing – original draft. **Michał Wawrzynowicz:** Conceptualization, Visualisation, Writing – original draft. **Katarzyna Markowska**: Data curation, Software. **Tomasz Chodkiewicz:** Data curation, Resources, Writing - review and editing. **Simon Butler:** Methodology, Validation, Writing - review and editing. **Lechosław Kuczyński:** Conceptualization, Data curation, Formal analysis, Funding acquisition, Methodology, Software, Supervision, Visualization, Writing – original draft.

## Acknowledgements

The study was supported by the National Science Centre (NCN) in Poland (grant no. 2018/29/B/NZ8/00066). The computational resources used in this work were provided by the Poznań Supercomputing and Networking Centre (grant no. pl0090-01). We are grateful to Klaudia Szala who tested the early versions of the models. The authors gratefully acknowledge MPPL volunteers for collecting the data, Chief Inspectorate for Environmental Protection (GIOŚ) and OTOP BirdLife Poland for providing the bird data.

## Declaration of generative AI and AI-assisted technologies in the manuscript preparation process

Artificial intelligence tools such as DeepL, ChatGPT and Gemini were used for language editing to improve grammar, clarity and readability of the manuscript. All the scientific content, including statistical analyses, results and conclusions was developed by the authors, who take full responsibility for the manuscript.

## Supporting Information

**Appendix A**. **Supplementary data.** Specific details concerning data used in the analysis, data treatment and variable selection.

**Appendix B. Supplementary data.** Additional results not included in the main article.

## Appendix A

### Bird abundance data

The analyses were carried out using real bird monitoring data for 14 grouped into 10 case studies. The dataset originates from the Common Breeding Bird Survey in Poland (MPPL) and spans 20 years (2001-2020), covering the entire area of Poland (more than 300 000 km^2^). The MPPL is a bird monitoring programme that is a part of The Pan-European Common Bird Monitoring Scheme (Brlík et al., 2021). Fieldworks in Poland began in 2000. Data collected between 2000 and 2006 were coordinated by OTOP BirdLife Poland, while from 2007 onward the surveys were conducted by OTOP under a contract with Chief Inspectorate for Environmental Protection (GIOŚ). The raw data for the analyses were obtained from both institutions. The main objective of the programme is to estimate population trends of the most common bird species. The fieldwork methodology is as follows: survey plots of 1 km^2^ are selected independently by stratified random sampling in 15 avifaunal regions of the country. Qualified volunteers visit survey plots twice during the breeding season (between 10 April and 15 May and between 16 May and 30 June) to conduct the bird counts along two 1-km long line transects separated by at least 0.5 km. During each visit, observers record all bird species seen or heard, with the distance band at which they were detected. The species densities (measured in pairs/km2) are then estimated using the distance sampling method (Buckland et al., 2001, 2004).

### Selected species

For the analysis we have selected pairs of species (Table A.1) that interact with each other through a mechanism **called social information transfer**, which relies on heterospecific attraction of focal species (cue users) to interacting species (cue providers) (Szymkowiak, 2013). These interactions can affect local populations in various ways, but we focused only on those that result in a local increase in the abundance of focal species in response to cues from interacting species. Each of the selected pairs exchanging social information was examined and confirmed experimentally (Table A.1). The density of the interacting species was included as an independent variable in models.

**Table A.1.** Bird species pairs selected for analysis.

| No. | Focal species (cue user) | Interacting species (cue provider) | Citation |
| --- | --- | --- | --- |
| 1. | Wood Warbler<br>(*Phylloscopus sibilatrix*) | Chiffchaff<br>(*Phylloscopus collybita*) | (Szymkowiak et al., 2017) |
| 2. | Pied flycatcher<br>(*Ficedula hypoleuca*) | Great Tit<br>(*Parus major*) | (Forsman et al., 1998; Mönkkönen & Forsman, 2002) |
| 3. | Blackcap<br>(*Sylvia atricapilla*) | Great Tit<br>(*Parus major*) | (Forsman et al., 2009) |
| 4. | Lesser Whitethroat<br>(*Curruca curruca*) | Great Tit<br>(*Parus major*) | (Forsman et al., 2009) |
| 5. | Garden Warbler<br>(*Sylvia borin*) | Blue Tit<br>(*Cyanistes caeruleus*) | (Forsman et al., 2009; Thomson et al., 2003) |
| 6. | Whitethroat<br>(*Curruca communis*) | Blue Tit<br>(*Cyanistes caeruleus*) | (Forsman et al., 2009) |
| 7. | Chaffinch<br>(*Fringilla coelebs*) | Willow Tit<br>(*Poecile montanus*) | (Forsman et al., 2009; Mönkkönen et al., 1990; Thomson et al., 2003) |
| 8. | Willow Warbler<br>(*Phylloscopus trochilus*) | Great Tit<br>(*Parus major*) | (Forsman et al., 1998, 2009; Mönkkönen & Forsman, 2002) |
| 9. | Common Redstart<br>(*Phoenicurus phoenicurus*) | Great Tit<br>(*Parus major*) | (Forsman et al., 1998) |
| 10. | Robin<br>(*Erithacus rubecula*) | Blue Tit<br>(*Cyanistes caeruleus*) | (Forsman et al., 2009; Thomson et al., 2003) |

### Environmental data

Based on published information about focal species, we selected two types of relevant dynamic environmental information: land-cover and climate. The data were imported into R using the ‘raster’ package (Hijmans, 2025), spatially queried to cover only Poland, temporally queried to match the study period (2001-2020), and reprojected to the Polish CS92 coordinate system (EPSG:2180) at a spatial resolution of 1 km. In addition, to account for the possible influence of past environmental conditions on current population densities, all dynamic variables were calculated with a one-year lag.

Land cover variables were obtained from ESA CCI Land Cover Maps v2.0.7 and v.2.11 (Defourny et al., 2017, 2024) provided by the Copernicus Climate Change Service. These variables represent the percentage cover of a given land cover class within each 300m × 300m grid cell in each year. Lagged versions of the land cover variables were highly collinear and were therefore excluded from the analyses.

Bioclimatic variables were extracted from TerraClimate, in a format of monthly global climate information with a spatial resolution of ∼4 km (Abatzoglou et al., 2018). For the study, we used minimum and maximum temperatures and precipitation. Due to the high collinearity of monthly climate data, they were aggregated into 3-month seasons (winter: December-February, spring: March-May, summer: June-August, autumn: September-November) by calculating their means (for temperatures) or sums (for precipitation).

Multicollinearity among predictors was assessed using pairwise correlations. From each pair of highly correlated variables, only one was retained, based on interpretability and ecological relevance. The final set of 19 environmental predictors was used in all case studies for each focal species (Table A.2).

**Table A.2.** List of predictor variables used in the analyses.

| No. | Variable name | Original name | Description | Included in the analysis |
| --- | --- | --- | --- | --- |
| Land cover variables from **ESA CCI Land Cover** |  |  |  |  |
| 1. | Rainfed_crops | 10 | rainfed crops [%] | No; collinearity |
| 2. | Herbaceous_crops | 11 | rainfed herbaceous crops [%] | No; collinearity |
| 3. | Tree_crops | 12 | rainfed shrub and tree crops [%] | No; too few levels |
| 4. | Mosaic_cropland | 30 | mosaic cropland (>50%) / natural vegetation (<50%) [%] | No; collinearity |
| 5. | Natural_vegetation | 40 | mosaic natural vegetation (>50%) / cropland (<50%) [%] | **Yes** |
| 6. | Deciduous_forest | 60 | broadleaved and deciduous forests closed to open (>15%) [%] | **Yes** |
| 7. | Coniferous_forest | 70 | needleleaved and evergreen forests closed to open (>15%) [%] | **Yes** |
| 8. | Mixed_forest | 90 | forests with mixed leaf type [%] | **Yes** |
| 9. | Shrub_herbaceous | 100 | mosaic tree and shrub cover (>50%) / herbaceous cover (<50%) [%] | **Yes** |
| 10. | Grassland | 130 | grassland [%] | **Yes** |
| 11. | Urban | 190 | urban areas [%] | **Yes** |
| 12. | Water | 210 | water bodies [%] | **Yes** |
| Climatic variables from **TerraClimate** |  |  |  |  |
| 13. | Precip_spring | ppt_3, ppt_4 and ppt_5 | logarithm of total precipitation in March, April and May [mm] | **Yes** |
| 14. | Precip_summer | ppt_6, ppt_7 and ppt_8 | logarithm of total precipitation in June, July and August [mm] | No; represent conditions after birds counting |
| 15. | Precip_autumn | ppt_9, ppt_10 and ppt_11 | logarithm of total precipitation in September, October and November [mm] | No; represent conditions after birds counting |
| 16. | Precip_winter | ppt_1, ppt_2 and ppt_12 | logarithm of total precipitation in December from previous year and January and February from current year [mm] | **Yes** |
| 17. | Tmax_spring | tmax_3, tmax_4 and tmax_5 | average of maximum temperature in March, April and May [°C] | No; collinearity |
| 18. | Tmax_summer | tmax_6, tmax_7 and tmax_8 | average of maximum temperature in June, July and August [°C] | No; represent conditions after birds counting |
| 19. | Tmax_autumn | tmax_9, tmax_10 and tmax_11 | average of maximum temperature in September, October and November [°C] | No; represent conditions after birds counting |
| 20. | Tmax_winter | tmax_1, tmax_2 and tmax_12 | average of maximum temperature in December from previous year and January and February from current year [°C] | No; collinearity |
| 21. | Tmin_spring | tmin_3, tmin_4 and tmin_5 | average of minimum temperature in March, April and May [°C] | **Yes** |
| 22. | Tmin_summer | tmin_6, tmin_7 and tmin_8 | average of minimum temperature in June, July and August [°C] | No; represent conditions after birds counting |
| 23. | Tmin_autumn | tmin_9, tmin_10 and tmin_11 | average of minimum temperature in September, October and November [°C] | No; represent conditions after birds counting |
| 24. | Tmin_winter | tmin_1, tmin_2 and tmin_12 | average of minimum temperature in December from previous year and January and February from current year [°C] | **Yes** |
| 25. | Precip_spring_lag | ppt_3, ppt_4 and ppt_5 | logarithm of total precipitation in March, April and May from previous year [mm] | **Yes** |
| 26. | Precip_summer_lag | ppt_6, ppt_7 and ppt_8 | logarithm of total precipitation in June, July and August from previous year [mm] | **Yes** |
| 27. | Precip_autumn_lag | ppt_9, ppt_10 and ppt_11 | logarithm of total precipitation in September, October and November from previous year [mm] | **Yes** |
| 28. | Precip_winter_lag | ppt_1, ppt_2 and ppt_12 | logarithm of total precipitation in December two years ago and January and February from previous year [mm] | No; too distant in the past |
| 29. | Tmax_spring_lag | tmax_3, tmax_4 and tmax_5 | average of maximum temperature in March, April and May from previous year [°C] | No; collinearity |
| 30. | Tmax_summer_lag | tmax_6, tmax_7 and tmax_8 | average of maximum temperature in June, July and August from previous year [°C] | **Yes** |
| 31. | Tmax_autumn_lag | tmax_9, tmax_10 and tmax_11 | average of maximum temperature in September, October and November from previous year [°C] | **Yes** |
| 32. | Tmax_winter_lag | tmax_1, tmax_2 and tmax_12 | average of maximum temperature in December two years ago and January and February from previous year [°C] | No; too distant in the past |
| 33. | Tmin_spring_lag | tmin_3, tmin_4 and tmin_5 | average of minimum temperature in March, April and May from previous year [°C] | **Yes** |
| 34. | Tmin_summer_lag | tmin_6, tmin_7 and tmin_8 | average of minimum temperature in June, July and August from previous year [°C] | **No** |
| 35. | Tmin_autumn_lag | tmin_9, tmin_10 and tmin_11 | average of minimum temperature in September, October and November from previous year [°C] | **Yes** |
| 36. | Tmin_winter_lag | tmin_1, tmin_2 and tmin_12 | average of minimum temperature in December two years ago and January and February from previous year [°C] | No; too distant in the past |

## Appendix B

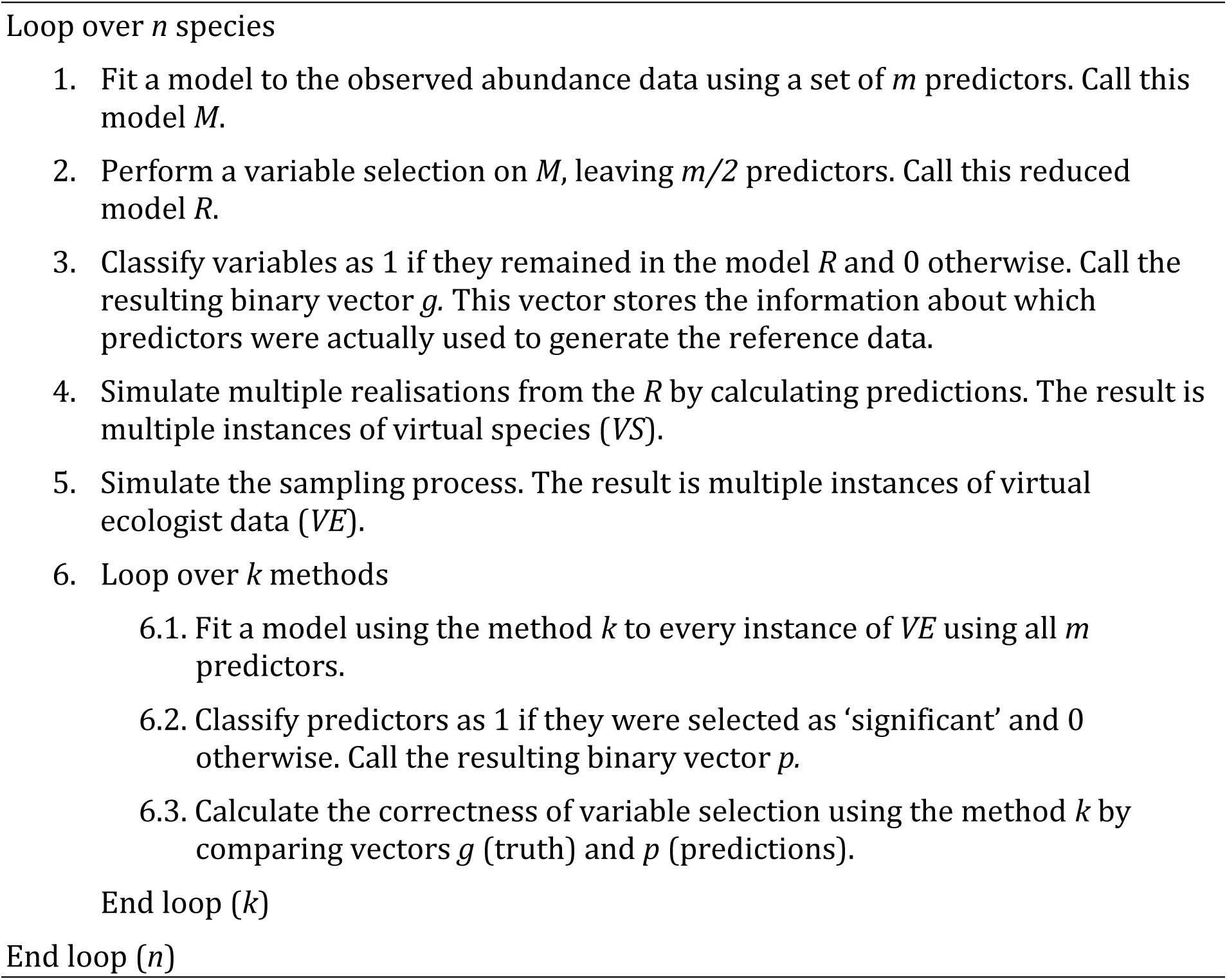
The algorithm used in the study.

**Table B.1.** Estimated marginal means of classification metrics: accuracy, true positive rate, and true negative rate for three compared approaches.

| Metric | Model | Without variable selection |  | With variable selection |  |
| --- | --- | --- | --- | --- | --- |
|  |  | Mean | 95% CI | Mean | 95% CI |
| Accuracy | TS | 0.50 | 0.48-0.53 | 0.55 | 0.52-0.57 |
|  | SDM | 0.68 | 0.66-0.70 | 0.63 | 0.61-0.65 |
|  | FRST | 0.88 | 0.87-0.89 | 0.77 | 0.76-0.79 |
| True positive rate (sensitivity) | TS | 0.00 | 0.00-1.00 | 0.14 | 0.11-0.17 |
|  | SDM | 0.68 | 0.62-0.73 | 0.73 | 0.68-0.78 |
|  | FRST | 0.84 | 0.81-0.88 | 0.84 | 0.81-0.88 |
| True negative rate (specificity) | TS | 1.00 | 0.00-1.00 | 0.95 | 0.95-0.96 |
|  | SDM | 0.69 | 0.66-0.72 | 0.54 | 0.51-0.57 |
|  | FRST | 0.93 | 0.92-0.94 | 0.71 | 0.68-0.74 |

---

ACCURACY

**Fig. B.1.**
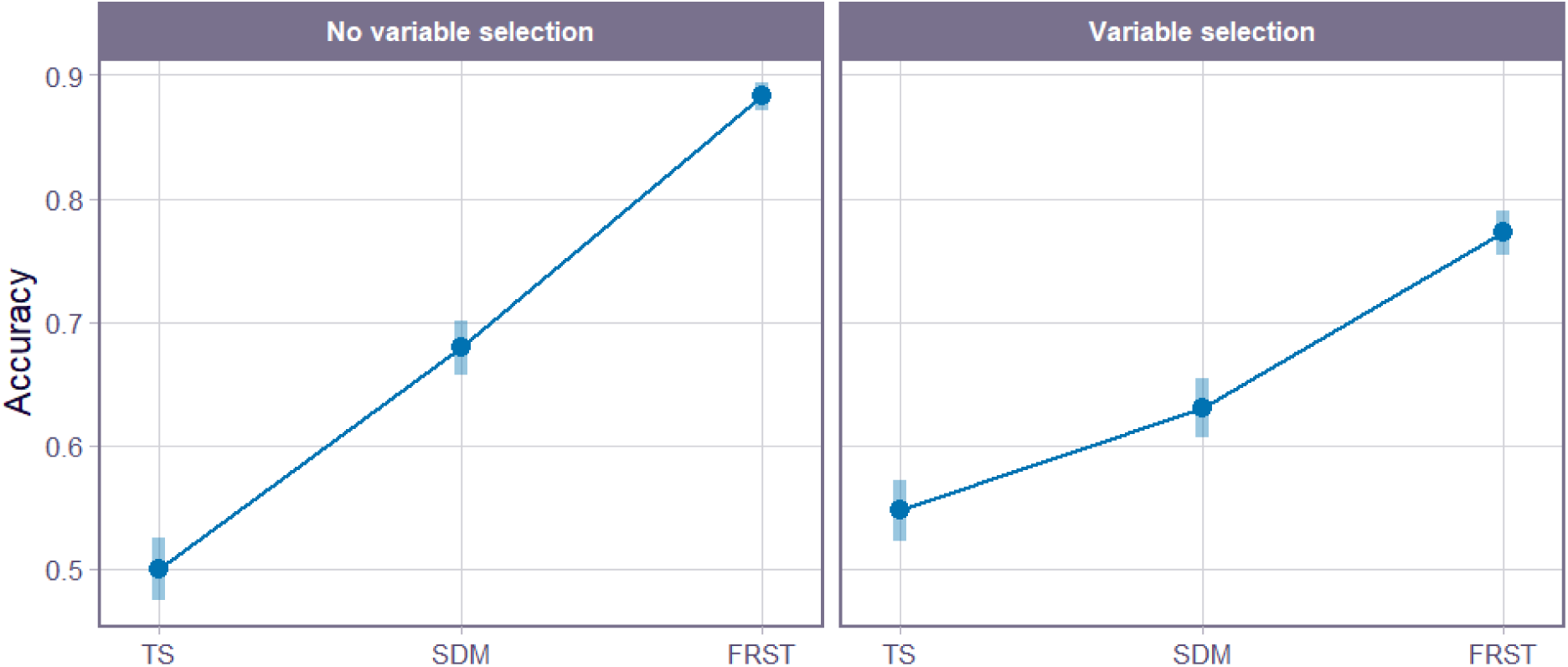
Effect of modelling approach (**TS**: time series, **SDM**: species distribution model, **FRST**: full-resolution spatio-temporal model) and variable selection (with or without variable selection) on model accuracy. Points represent estimated marginal means, blue bars indicate 95% confidence intervals.

**Table B.2.** Estimated differences in marginal means between variable selection options for each modelling approach.

| Model | Contrast<br>(variable selection) | odds.ratio | SE | z.ratio | p.value |
| --- | --- | --- | --- | --- | --- |
| TS | no / yes | 0.83 | 0.02 | -9.49 | <.0001 |
| SDM | no / yes | 1.24 | 0.03 | 10.28 | <.0001 |
| FRST | no / yes | 2.22 | 0.06 | 28.74 | <.0001 |

**Table B.3.**
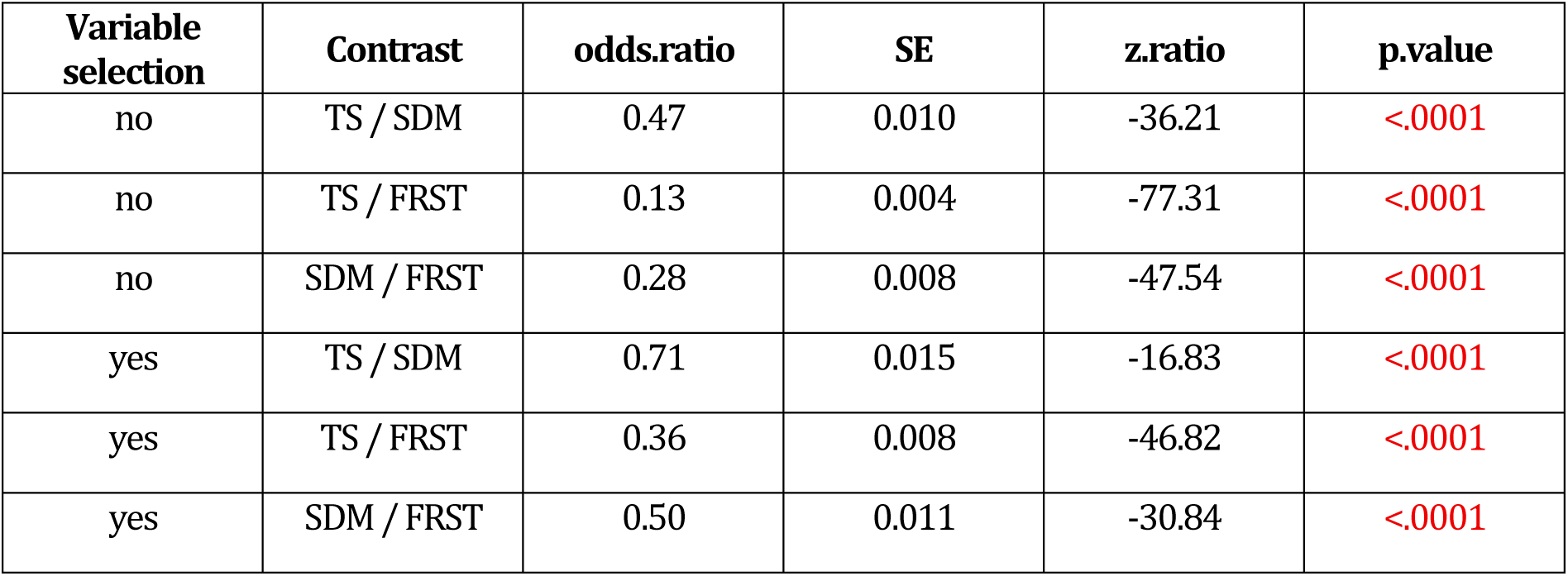
Estimated differences in marginal means between different modelling approaches for each variable selection option.

TRUE POSITIVE RATE / SENSITIVITY

**Fig. B.2.**
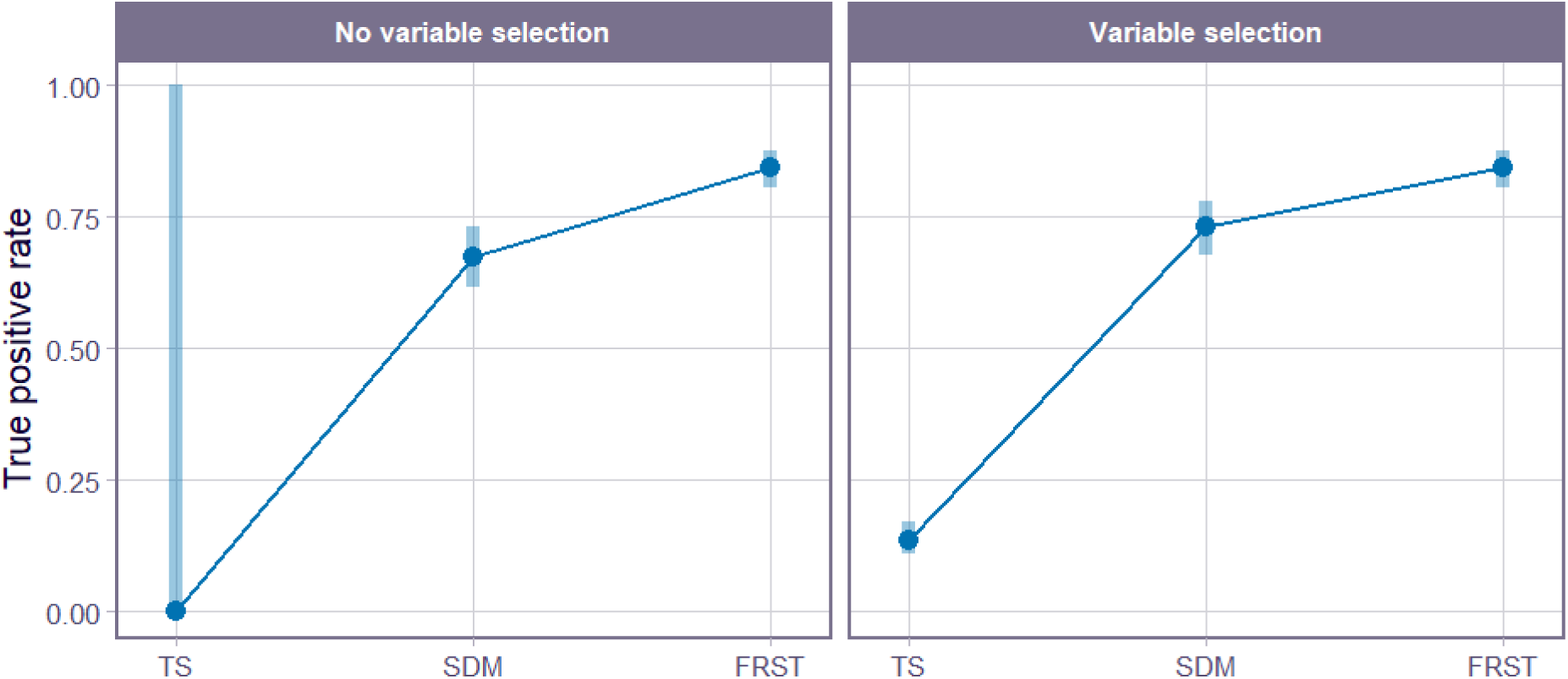
Effect of modelling approach (**TS**: time series, **SDM**: species distribution model, **FRST**: full-resolution spatio-temporal model) and variable selection (with or without variable selection) on model sensitivity. Points represent estimated marginal means, blue bars indicate 95% confidence intervals.

**Table B.4.**
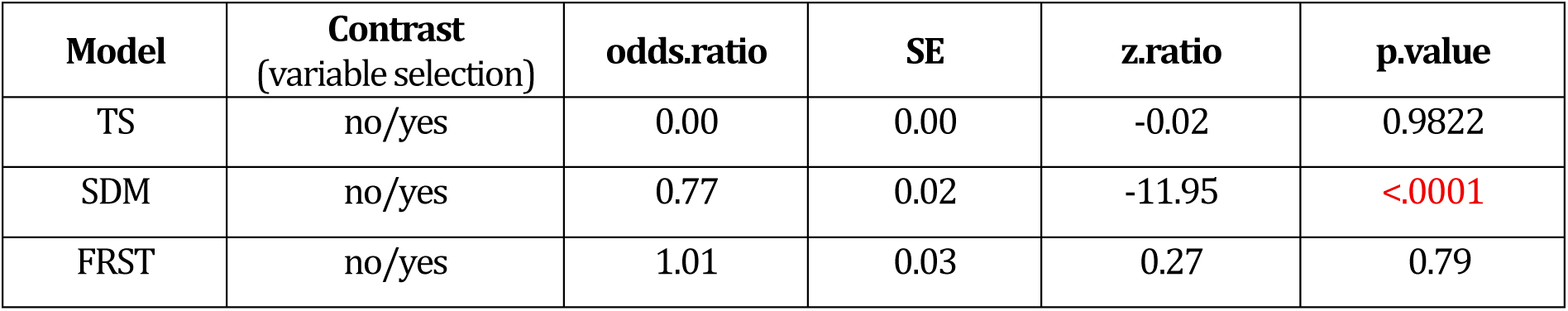
Estimated differences in marginal means between variable selection options for each modelling approach.

**Table B.5.** Estimated differences in marginal means between different modelling approaches for each variable selection option.

| Variable selection | Contrast | odds.ratio | SE | z.ratio | p.value |
| --- | --- | --- | --- | --- | --- |
| no | TS / SDM | 0.00 | 0.00 | -0.03 | 0.9997 |
| no | TS / FRST | 0.00 | 0.00 | -0.03 | 0.9996 |
| no | SDM / FRST | 0.38 | 0.01 | -38.83 | <.0001 |
| yes | TS / SDM | 0.06 | 0.00 | -108.45 | <.0001 |
| yes | TS / FRST | 0.03 | 0.00 | -124.19 | <.0001 |
| yes | SDM / FRST | 0.50 | 0.01 | -27.31 | <.0001 |

TRUE NEGATIVE RATE / SPECIFICITY

**Fig. B.3.**
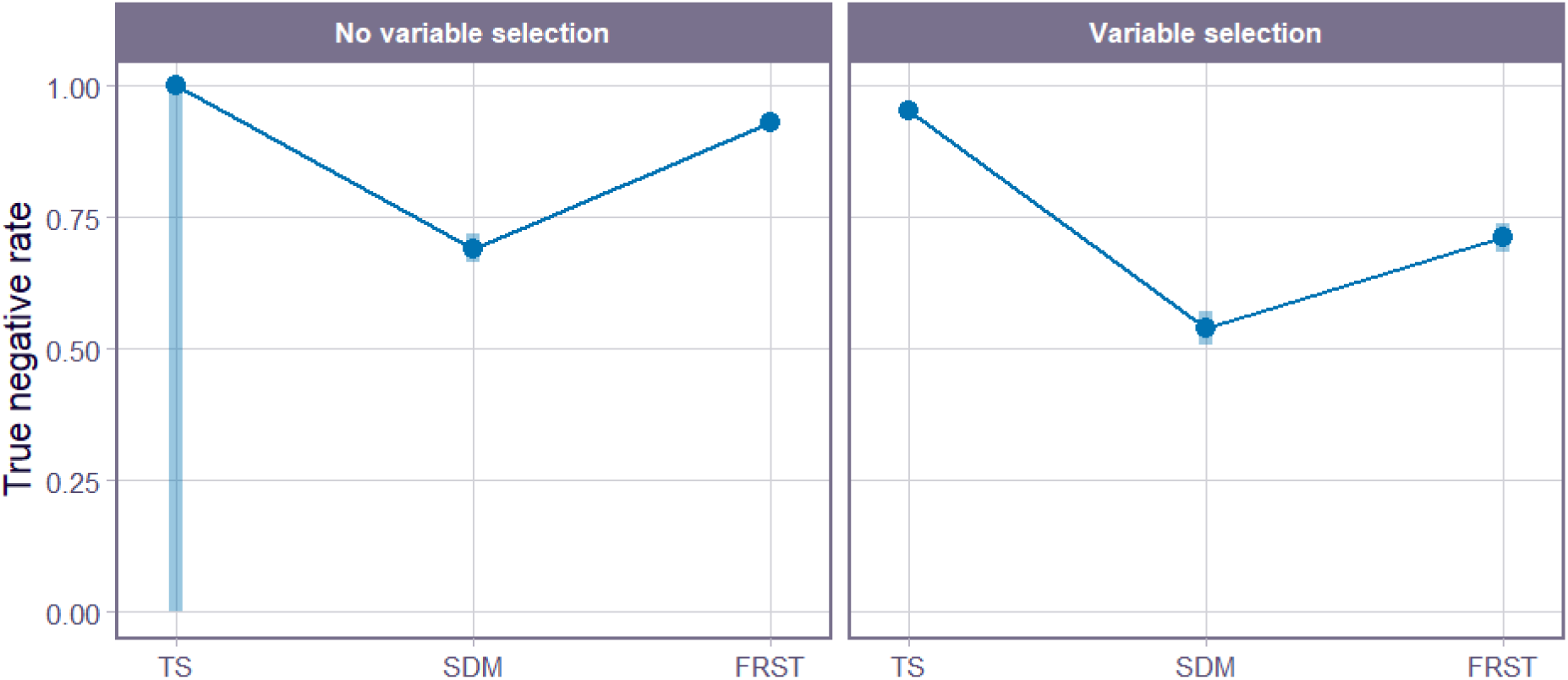
Effect of modelling approach (**TS**: time series, **SDM**: species distribution model, **FRST**: full-resolution spatio-temporal model) and variable selection (with or without variable selection) on model specificity. Points represent estimated marginal means, blue bars indicate 95% confidence intervals.

**Table B.6.** Estimated differences in marginal means between variable selection options for each modelling approach.

| Model | Contrast (variable selection) | odds.ratio | SE | z.ratio | p.value |
| --- | --- | --- | --- | --- | --- |
| TS | no/yes | 8.3e+08 | 7.45e+11 | 0.023 | 0.9817 |
| SDM | no/yes | 2.00 | 0 | 31.115 | <.0001 |
| FRST | no/yes | 5.00 | 0 | 53.152 | <.0001 |

**Table B.7.** Estimated differences in marginal means between different modelling approaches for each variable selection option.

| Variable selection | Contrast | odds.ratio | SE | z.ratio | p.value |
| --- | --- | --- | --- | --- | --- |
| no | TS / SDM | 7.37e+09 | 6.61e+12 | 0.03 | 0.9996 |
| no | TS / FRST | 1.25e+09 | 1.13e+12 | 0.02 | 0.9997 |
| no | SDM / FRST | 0.00 | 0.00 | -56.38 | <.0001 |
| yes | TS / SDM | 17.00 | 1.00 | 79.17 | <.0001 |
| yes | TS / FRST | 8.00 | 0.00 | 57.46 | <.0001 |
| yes | SDM / FRST | 0.00 | 0.00 | -35.25 | <.0001 |

**Figure B.4.**
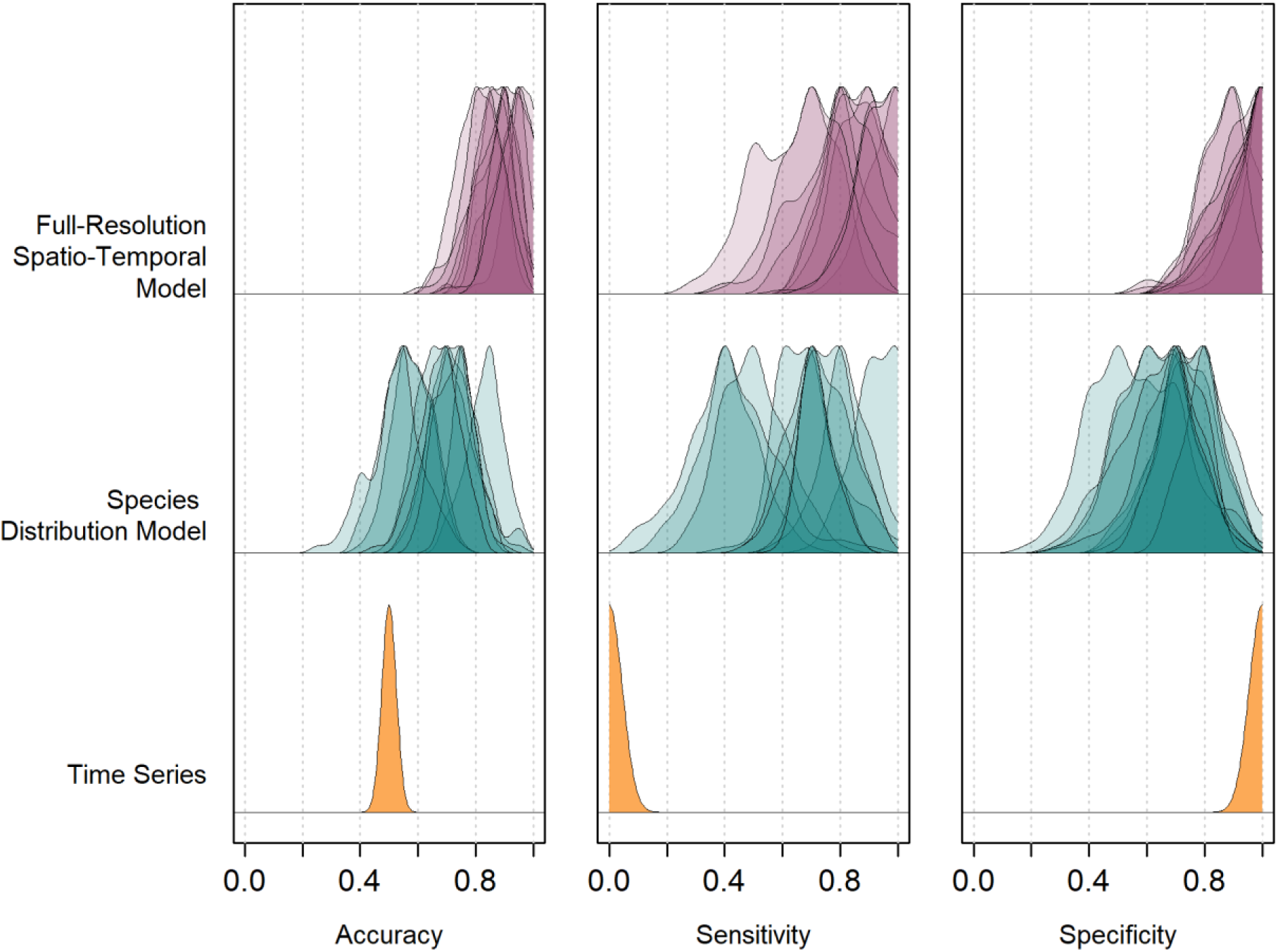
Distributions of model performance metrics (accuracy, sensitivity and specificity) for three modelling approaches (time series, species distribution model and full-resolution spatio-temporal model). Kernel density curves represent values obtained for individual case studies. No variable selection method was applied to the final models.

**Figure B.5.**
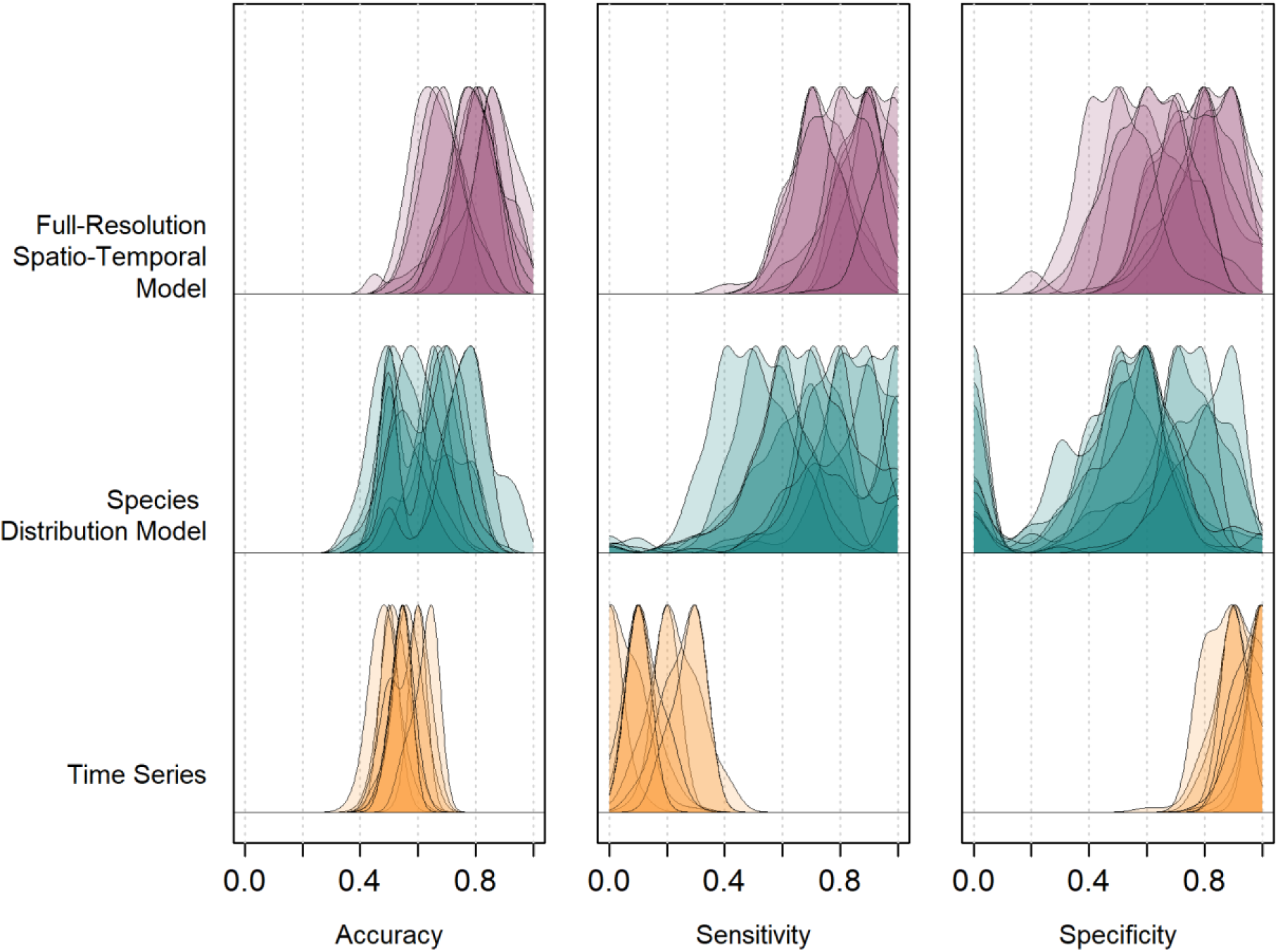
Distributions of model performance metrics (accuracy, sensitivity and specificity) for three modelling approaches (time series, species distribution model and full-resolution spatio-temporal model). Kernel density curves represent values obtained for individual case studies. Double penalty shrinkage was applied to the final models.

**Figure B.6.**
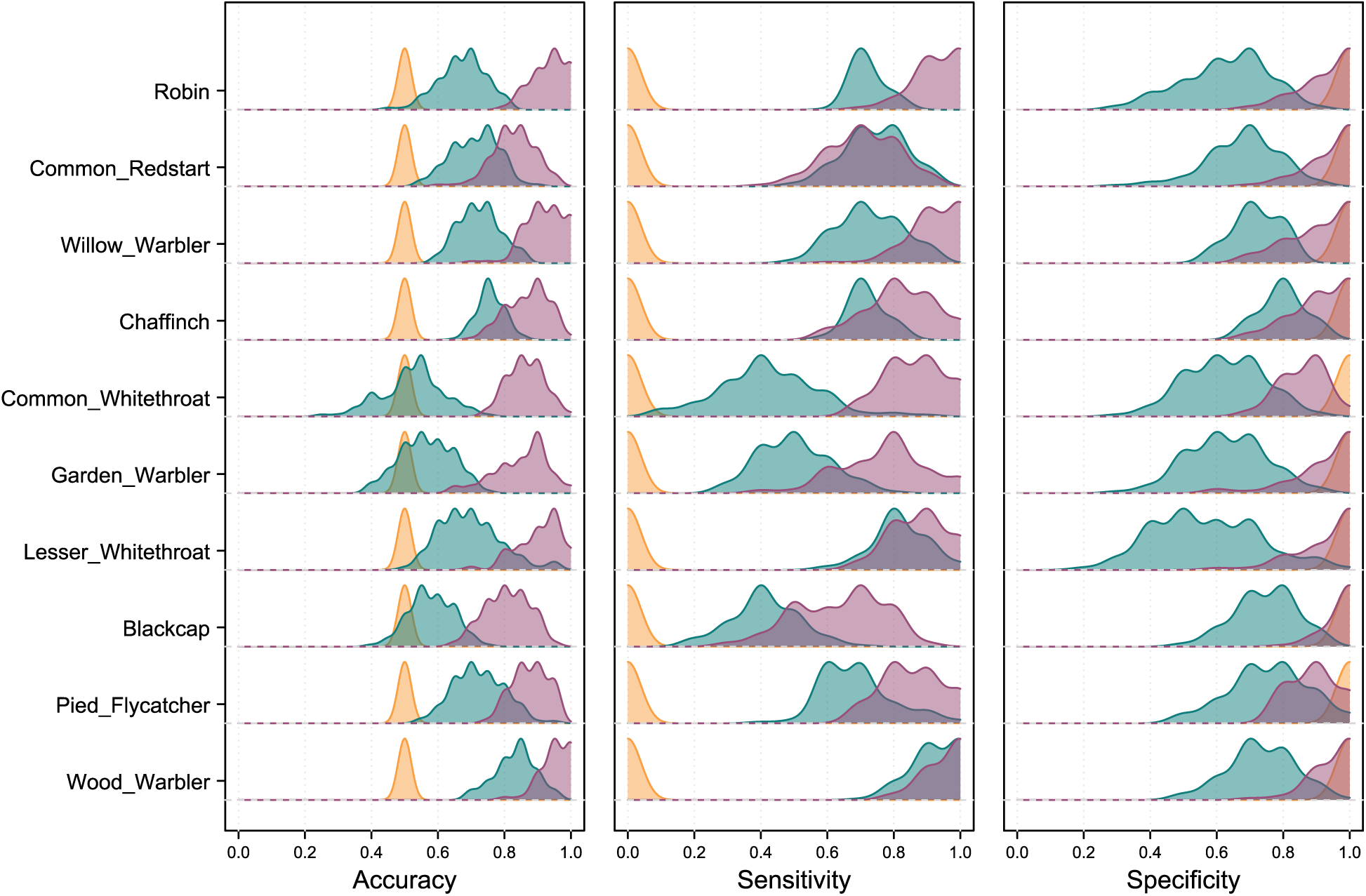
Distributions of model performance metrics (accuracy, sensitivity, and specificity) across individual case studies (bird species). The colour of each density curve indicates the modelling approach used; yellow: time series (TS), green: species distribution model (SDM), and purple: full-resolution spatio-temporal (FRST) model. No variable selection was applied to the final models.

**Figure B.7.**
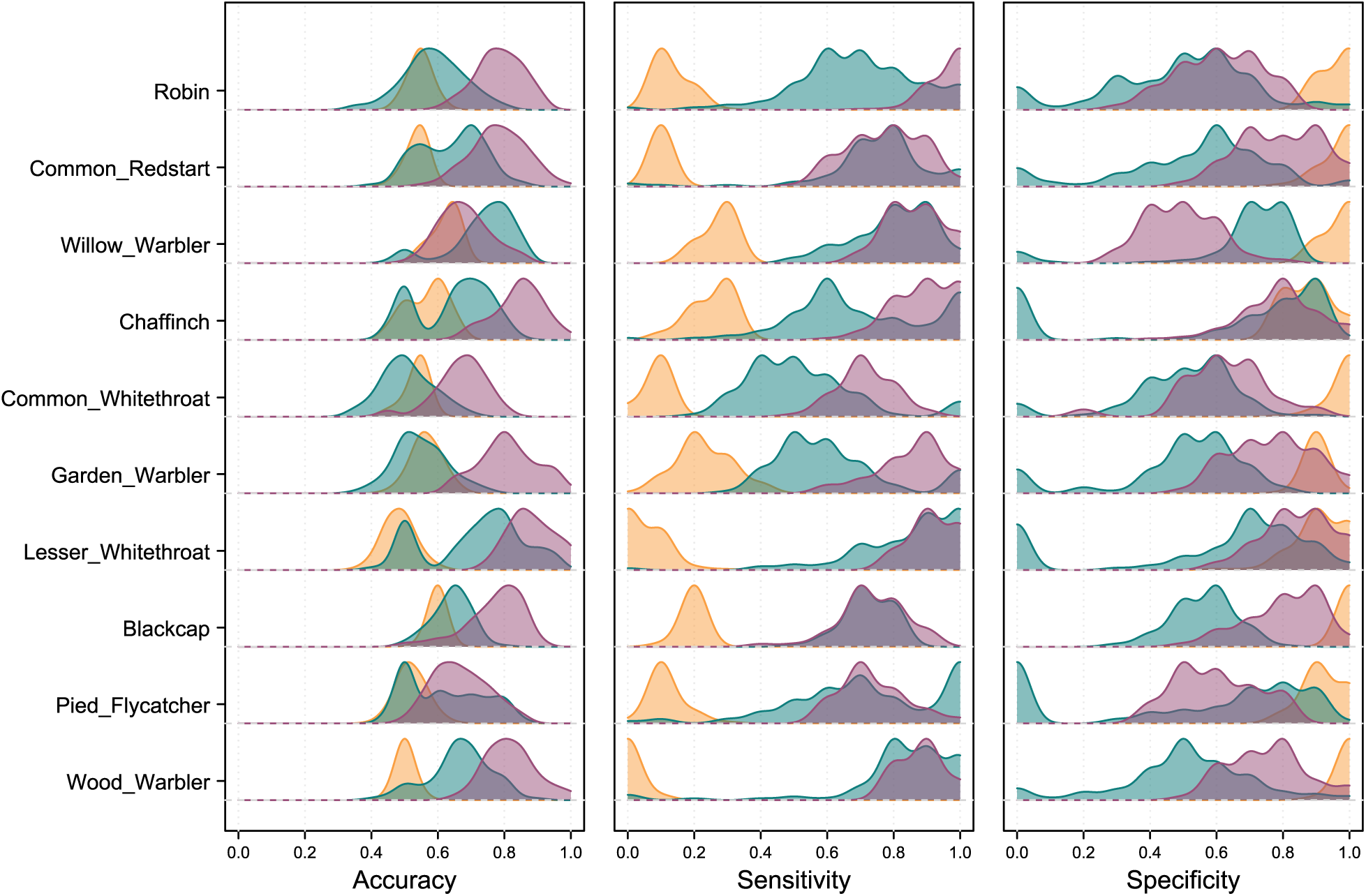
Distributions of model performance metrics (accuracy, sensitivity and specificity) across individual case studies (bird species). The colour of each density curve indicates the modelling approach used; yellow: time series (TS), green: species distribution model (SDM), and purple: full-resolution spatio-temporal (FRST) model. Double penalty shrinkage was applied to the final models.

